# Bacteriophage Kil peptide folds into a predicted helix-turn-helix structure to disrupt *Escherichia coli* cell division

**DOI:** 10.1101/2024.12.16.628577

**Authors:** Arindam Naha, Todd A. Cameron, William Margolin

## Abstract

FtsZ, a eukaryotic tubulin homolog and an essential component of the bacterial divisome, is the target of numerous antimicrobial compounds as well as proteins and peptides, most of which inhibit FtsZ polymerization dynamics. We previously showed that the Kil peptide from bacteriophage λ inhibits *Escherichia coli* cell division by disrupting FtsZ ring assembly, and this inhibition requires the presence of the essential FtsZ membrane anchor protein ZipA. To investigate Kil’s molecular mechanism further, we employed truncation mutants and molecular modeling to identify the minimal residues necessary for its activity. Modeling suggests that Kil’s core segment folds into a helix-turn-helix (HTH) structure. Deleting either the C-terminal 11 residues or the N-terminal 5 residues of Kil still allowed inhibition of *E. coli* cell division, but removing both termini nearly abolished this activity, indicating that a minimal region within the Kil HTH core is essential for its function. Another Kil-like peptide from a closely related enterobacterial phage also disrupts FtsZ ring assembly and requires ZipA for this activity. Consistent with its broader activity against FtsZ, λ Kil was able to efficiently inhibit cell division of a uropathogenic *E. coli* (UPEC) strain. Understanding the function of Kil and similar peptides can potentially reveal additional ways to target FtsZ for antimicrobial therapies as well as elucidate how FtsZ functions in bacterial cell division.

## 1. Introduction

One of the most formidable challenges a bacterial cell faces during survival and proliferation is to faithfully duplicate and partition its genome and then divide into two identical daughter cells, a process known as cytokinesis [1,2]. Binary fission, the primary method of bacterial cell division, is driven by a multiprotein complex called the divisome. The formation of the divisome begins with the polymerization of FtsZ monomers into protofilaments, resulting in the formation of a ring-like structure, known as the Z-ring, at the cell’s midpoint [3,4]. This Z-ring forms at the site of future division, marking the first crucial step in bacterial cytokinesis. Following Z-ring assembly, the remaining divisome proteins are recruited through multiple interactions [5,6].

Proper localization and recruitment of the Z-ring is critical and intricately regulated. The assembly and function of the Z-ring during cell division relies on the interaction of FtsZ polymers with the cytoplasmic membrane (CM). FtsZ lacks intrinsic membrane-binding ability, necessitating the involvement of adaptor proteins that bridge FtsZ and the membrane [7]. These anchors not only attach FtsZ to the membrane but also organize it into higher-order structures that enhance cell division. Key membrane tethers, such as FtsA and SepF, are conserved across diverse bacterial species [7]. They attach to the membrane using an amphipathic helix and form various oligomeric structures [8–11]. In contrast, less-conserved proteins like EzrA and ZipA possess transmembrane domains and interact with FtsZ through multiple binding sites, forming extended structures [12–15].

In γ-proteobacteria like *Escherichia coli*, both FtsA and ZipA are essential for cell division. These proteins independently tether FtsZ polymers to the CM. While either FtsA or ZipA can localize Z rings to midcell, both are required for functional divisome assembly and septum formation [16]. The absence of both proteins prevents Z ring formation entirely, underscoring the necessity of at least one tether for FtsZ’s membrane attachment and Z ring structure.

Bacterial antibiotic resistance is a global threat to human health [17]. As bacteria quickly develop resistance, our arsenal of effective drugs is depleted, necessitating the discovery of new antibiotic targets. One very promising target is the bacterial divisome [18]. Due to its pivotal role in bacterial cell division and highly conserved sequence homology across prokaryotes [19], FtsZ, a bacterial homologue of tubulin and the divisome’s keystone, has emerged as an attractive target for antibiotic development [20]. Consistent with its key role, many naturally occurring small molecules and proteins are known to bind to FtsZ and perturb its essential function [21]. Inhibiting FtsZ activity results in the disruption of bacterial cytokinesis: rod-shaped cells such as *Escherichia coli* elongate into filaments and eventually lyse, resulting in total loss of viability (9).

For billions of years, bacteria and their phages (bacteriophages) have been engaged in an evolutionary arms race [22]. After several decades of study, we now know that some bacteriophage-encoded peptides interfere with host cell morphology or cytokinesis by targeting FtsZ during viral infection, leading to cell filamentation and death [23–26]. One of the best studied of these is Kil, a 47 amino acid peptide produced by bacteriophage lambda (λ) during lytic growth. It was previously shown that Kil inhibits *E. coli* cell division by perturbing Z-ring assembly, and this inhibition requires the essential FtsZ membrane tether ZipA [25,27]. In this study, as a first step to define Kil’s molecular mechanism of action, we have used molecular modeling and mutagenesis to define the minimal residues required for Kil’s cell division inhibition activity. We also tested the ability of a related Kil-like phage-encoded peptide to inhibit *E. coli* cell division, as well as the ability of λ Kil to inhibit cell division of uropathogenic *E. coli* (UPEC).

## 2. Results

### 2A. Kil requires a helix-turn-helix structural element for its anti-FtsZ activity

Modeling by AlphaFold 3 [28] indicated that the His_6_-tagged λ Kil peptide is predicted to form a helix-turn-helix (HTH) structure (Fig. 1, top). The N-terminal (aa 4-18) and C-terminal (aa 21-38) domains were predicted with very high confidence (predicted local distance difference test, pLDDT, value >90) to form the two arms of the HTH structure, helix 1 and 2, respectively. Both helices are interlinked by a turn comprising a glycine and aspartic acid (G19-D20).

**Figure 1.**
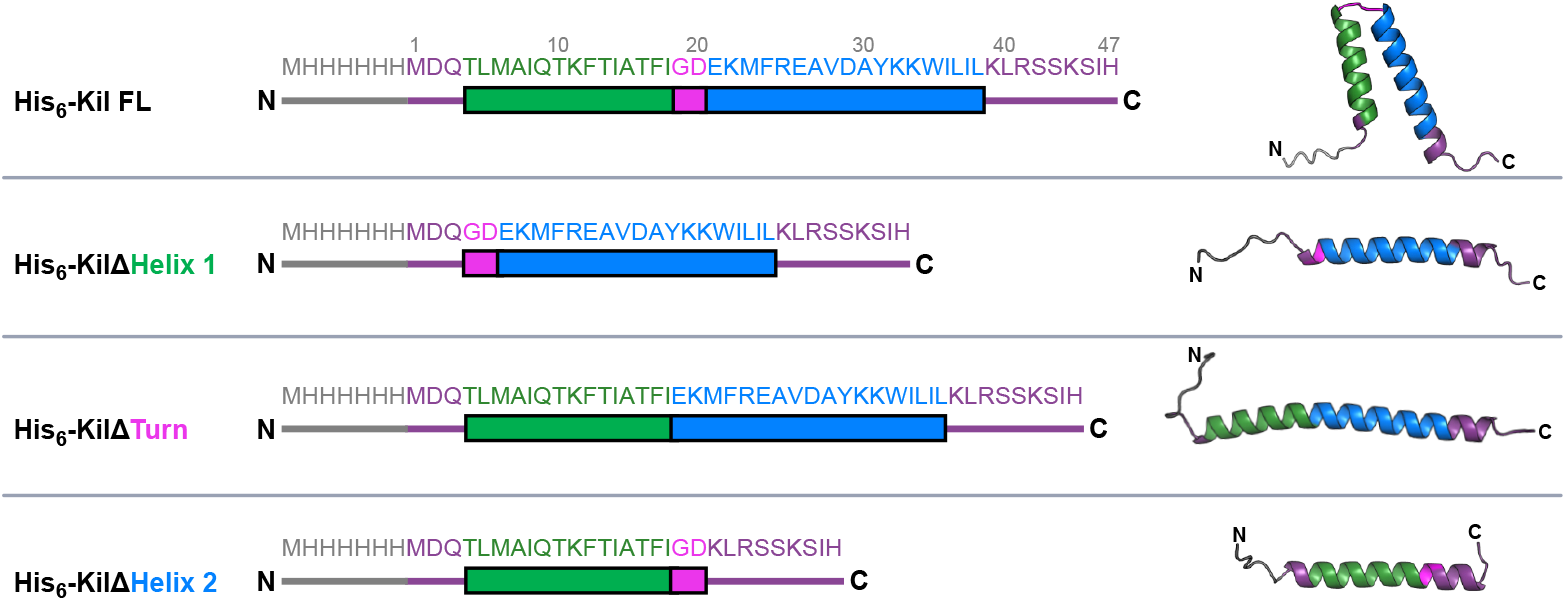
Schematic representation of the λ Kil peptide and deletion derivatives. Green and blue bars indicate helix 1 and helix 2, respectively, with the turn highlighted in magenta and the 6xHis tag highlighted in grey. The full length (FL) Kil peptide with its intact HTH domains is depicted in the top row, followed by derivatives deleted for helix 1, the turn, or helix 2, shown below. Amino acid numbering of the native Kil peptide is shown above the FL sequence. Ribbon diagrams of each His-tagged Kil derivative are shown as high confidence predictions from AlphaFold 3, with corresponding colors representing the different domains.

To acquire more details about the functional importance of this HTH structure in retaining Kil’s anti-FtsZ activity, we generated N-terminally His_6_-tagged Kil mutants in which the residues comprising helix 1, helix 2, or the inter-helix turn were deleted. As expected, when these HTH mutants were modeled using AlphaFold 3 (Fig. 1), the resulting structures lacked Kil’s signature HTH core. We then investigated the ability of these His-tagged Kil derivatives to inhibit cytokinesis in the *E. coli* K-12 strain WM1074 when expressed from the arabinose-inducible *araBAD* promoter on the multi-copy plasmid pBAD24.

Microscopic examination of WM1074 cells expressing full-length (FL) Kil or the truncated mutants demonstrated that an intact HTH core is indispensable for Kil’s activity (Fig. 2A). Under the repressing condition, where the *pBAD* promoter is shut off in the presence of glucose, the bacterial cells divided normally. Following arabinose induction, His-tagged FL-Kil produced from pBAD24 inhibited cytokinesis and formed filamentous cells as previously described [25]. In contrast, we observed a complete suppression of cell division inhibition in *E. coli* cells expressing either of the His-tagged Kil HTH truncations (Fig. 2A ΔHelix1, ΔHelix2 or ΔTurn).

**Figure 2.**
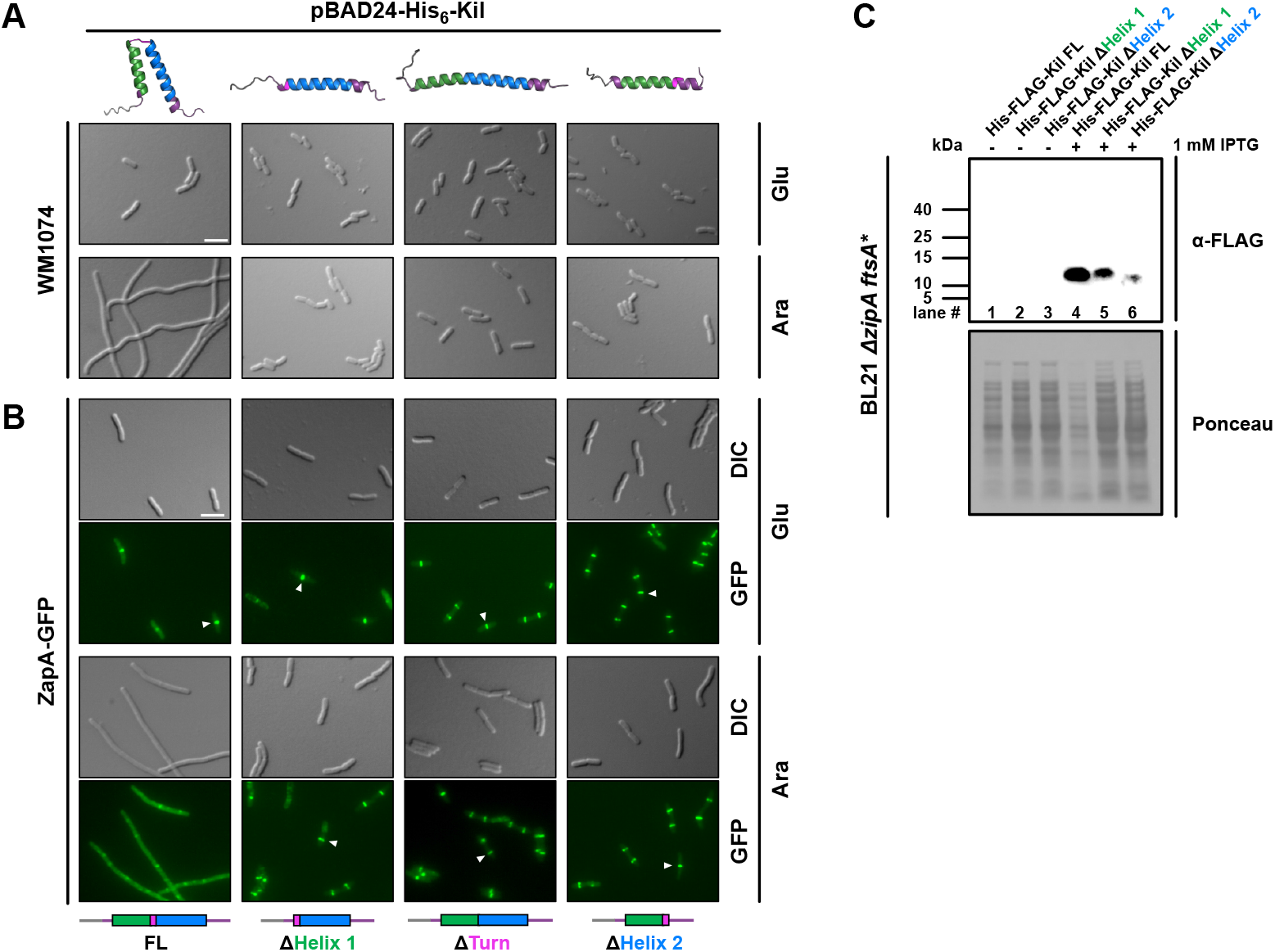
The ability of the λ Kil peptide to inhibit cell division requires an intact HTH core. (A) DIC (differential interference contrast) microscopic images of WT *E. coli* strain WM1074 expressing His-tagged FL Kil or its HTH truncations from pBAD24 under repressing (0.2% glucose, Glu) or inducing (0.2% arabinose, Ara) conditions. All derivatives are fused to an N-terminal His epitope. (A and B) Linear schematics and AlphaFold 3-modeled structures of the various derivatives are presented below and above each column of corresponding micrographs. Scale bars, 5 µm. (B) Fluorescence microscopic images of strain WM4952 expressing *zapA*-GFP plus either FL Kil or HTH deletion derivatives of Kil described in part A. Arrowheads highlight intact Z-rings. (C) Immunoblot of His-FLAG-FL Kil (lanes 1 and 4) and His-FLAG-Kil variants with either helix1 (lanes 2 and 5) or helix2 (lanes 3 and 6) deleted, expressed from a phage T7 promoter in pET15b ± IPTG induction in strain WM5017 (Δ*zipA ftsA\**) to suppress Kil toxicity, and probed with α-FLAG to detect expression of the tagged Kil derivatives. The predicted sizes of the His-FLAG-tagged FL, Δhelix 1 or Δhelix 2 Kil derivatives are 10.5, 8.8 and 8.2 kDa, respectively. A portion of the Ponceau S-stained blot is shown below the immunoblot as a loading control.

We further investigated whether the resistance of *E. coli* cells to Kil HTH truncations could be due to unperturbed Z-ring assembly, hypothesizing that HTH mutants might lose their ability to bind and/or inhibit FtsZ polymerization. To evaluate Z-ring localization, we used *E. coli* cells expressing ZapA-GFP at the native locus (WM4952) as a proxy, as ZapA strongly localizes to the divisome through direct interactions with FtsZ [29,30]. As expected, WM4952 cells expressing His-tagged full-length (FL) Kil exhibited ZapA-GFP dispersed throughout the filamentous cells, indicating Z-ring disintegration (Fig. 2B). In stark contrast, *E. coli* cells producing HTH variants exhibited intact Z rings even one hour after arabinose induction.

Despite repeated attempts, we were unable to detect His-tagged Kil derivatives in either *E. coli* WM1074 or WM4952 by immunoblotting. Therefore, to quantitatively measure intracellular accumulation levels of the Kil derivatives, we expressed His-FLAG-tagged FL Kil and two of the HTH mutants, ΔHelix1 and ΔHelix2, from the high-expression vector pET15b. We used a *ΔzipA* strain carrying the *ftsA*\* allele previously used for Kil purification [25,27] because Kil activity is ZipA-dependent and thus is less toxic to cells lacking ZipA. However, as ZipA is essential for *E. coli* cell division, to permit viability we combined *ΔzipA* with a gain-of-function allele in another FtsZ anchor protein, FtsA^R286W^ (*ftsA*\*) that bypasses the need for several normally essential cell division proteins, including ZipA and FtsK [31]. Notably, we found that both Kil truncations, particularly Kil ΔHelix2, were unstable in the cell lysates extracted from BL21 *ΔzipA ftsA*\* recombinant strains (Fig. 2C). Unfortunately, we were unable to generate a pET15b His-FLAG-KilΔTurn despite repeated efforts. Taken together, this result supports our speculation that an unperturbed HTH core is necessary for Kil’s stability and anti-FtsZ activity in *E. coli* cells.

### 2B. Identifying the minimal region of Kil that inhibits cell division

Given that residues 4-38 of Kil are predicted to form the HTH structure (Fig. 1), we hypothesized that truncations at the N-terminal and C-terminal ends of the 47-residue Kil sequence might retain functional activity. To test this hypothesis, we generated several deletions at the N-terminus and C-terminus of the His-tagged full-length (FL) Kil peptide, modeled their structures with AlphaFold 3 (Fig. 3A), and functionally expressed these variants. We then compared the ability of these truncated versions to induce filamentous cell formation, indicative of cell division inhibition, with that of the His-Kil FL peptide.

**Figure 3.**
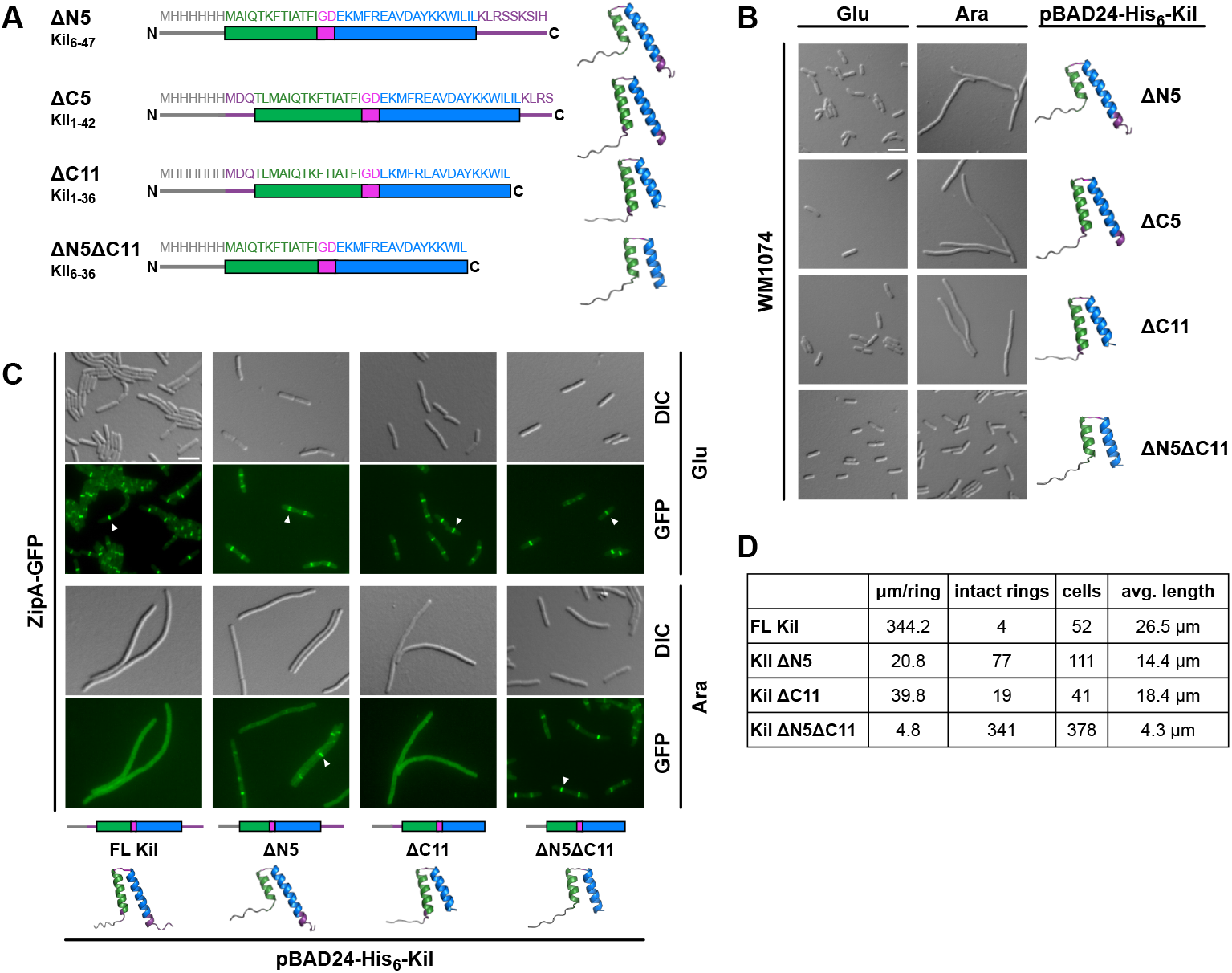
Shorter λ Kil truncations differentially inhibit *E. coli* cytokinesis. (A) Additional truncation derivatives of His-tagged λ Kil are diagrammed, using the same color schemes as in Fig. 1. (B) DIC microscopic images of WM1074 expressing His-tagged Kil truncations from pBAD24 under repressing (0.2% glucose, Glu) or inducing (0.2% arabinose, Ara) conditions. Scale bar, 5 µm. AlphaFold 3-modeled structures of the Kil derivatives are included. (C) Fluorescence microscopic images are shown of strain XTL937 expressing ZipA-GFP at its native chromosomal locus along with the indicated His-tagged Kil derivatives in pBAD24. Arrowheads indicate intact Z-rings. Schematic diagrams of the Kil derivatives are presented at the bottom of each corresponding micrograph, along with AlphaFold 3-modeled structures. Scale bar, 5 µm. (D) Quantitation of ZipA-GFP rings from the strains shown in panel C induced with arabinose.

Our experiments revealed that deletions of up to 11 amino acids from the C-terminus (KilΔC11, corresponding to residues 1–36) or the first 5 amino acids from the N-terminus (KilΔN5, corresponding to residues 6–47) did not impair Kil’s ability to inhibit *E. coli* cell division (Fig. 3B). However, simultaneous deletion of both the N-terminal five amino acids and the 11 C-terminal amino acids (KilΔN5ΔC11, corresponding to residues 6–36) completely abolished Kil activity, resulting in short, dividing cells (Fig. 3B). Using ZipA-GFP as a proxy for Z rings, we found that induction of KilΔC11 or KilΔN5 from pBAD24 disassembled most Z rings, although ∼10-fold less efficiently compared with His-FL Kil, whereas induction of KilΔN5ΔC11 left Z rings intact in all cells, as expected (Fig. 3C-D). Induction of KilΔN5 (1 intact Z ring per ∼20 µm of cell length) was slightly less effective at disassembling Z rings compared with KilΔC11 (1 intact Z ring per ∼44 µm of cell length) (Fig. 3C-D), but both were clearly potent at inhibiting Z ring formation. Similar patterns of behavior were observed after induction of the Kil derivatives in the WM4952 strain expressing ZapA-GFP (Fig. S1).

### 2C. Kil peptide from Enterobacteria phage HK629 shares a common mechanism to inhibit bacterial cytokinesis in *E. coli*

Kil peptides are diverse and widely distributed among enterobacterial lambda-like phages. For example, a 37 amino acid peptide encoded by Enterobacterial lambdoid phage HK629 shares significant sequence similarity with λ Kil, including most of the λ Kil HTH core (Fig. 4A). As a result, HK629 Kil closely resembles λ KilΔC11 except that HK629 Kil’s C terminus consists of the highly polar triplet RSK instead of the aliphatic residues that terminate λ KilΔC11. Curiously, a similar RSSK motif exists in the C-terminus of full length λ Kil, so we hypothesized that that RSK in HK629 Kil may be of functional importance. Notably, a five amino acid N-terminal truncation of HK629 Kil, which we term HK629 Kil’, is nearly identical in sequence to λ KilΔN5ΔC11 aside from the presence of the RSK triplet sequence.

**Figure 4.**
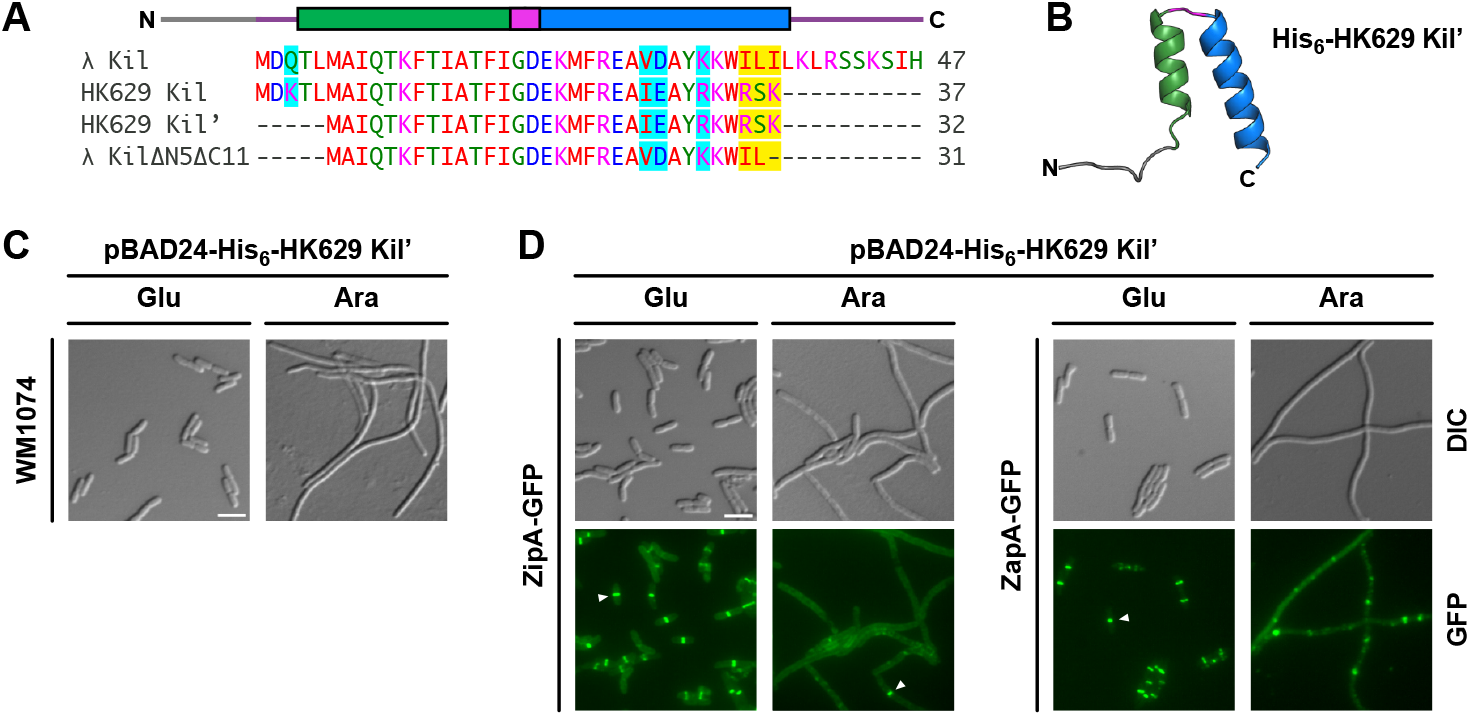
Comparing cell division inhibition activities of Kil from phage HK629 and λ. (A) Sequence alignment of Kil peptides produced by bacteriophages λ and HK629 using the Clustal Omega multiple sequence alignment tool. HK629 Kil has several potential initiator methionines; longer (Kil) and shorter (Kil’) versions are depicted. Similar amino acids are highlighted in cyan and non-conservative amino acid positions are highlighted in yellow. λ Kil HTH domains are depicted above the alignment. (B) Predicted structure of His-tagged HK629 Kil’ from AlphaFold 3, showing a HTH fold similar to λ Kil. (C) DIC microscopy of pBAD24-His-HK629-Kil’ expressed in WT strain WM1074. Scale bar, 5 µm. (D) DIC (top) or fluorescence (bottom) microscopy of pBAD24-His-HK629-Kil’ expressed in strain XTL937 with ZipA-GFP coexpressed from the native locus (left panels) or in strain WM4952 expressing chromosomal ZapA-GFP (right panels). Expression of His-HK629 Kil’ was either repressed (Glu) or induced (Ara), showing that HK629 Kil’ inhibits cell division and disrupts Z rings. Arrowheads highlight intact Z-rings. Scale bar, 5 µm.

Therefore, we set out to determine whether HK629 Kil’ would successfully inhibit bacterial cell division like λ Kil.

As expected, AlphaFold 3 predicted that a His-tagged 39-residue version of HK629 Kil’ can form a HTH structure similar to that of λ Kil (Fig. 4B). Next, we expressed the His-HK629 Kil’ (analogous to His-λ KilΔN5ΔC11) from pBAD24, with the hypothesis that this would act to inhibit *E. coli* cell division. Indeed, induction of His-HK629 Kil’ in *E. coli* WM1074 resulted in the inhibition of cytokinesis (Fig. 4C). As with λ Kil, HK629 Kil’ disrupted most Z rings, as measured by fluorescence microscopy of cells expressing HK629 Kil’ along with either ZipA-GFP or ZapA-GFP expressed from their native chromosomal loci as proxies for Z rings (Fig. 4D).

To investigate whether HK629 Kil’ requires ZipA to inhibit *E. coli* cell division like λ Kil, we expressed His-HK629 Kil’ in wild-type (WT) *zipA ftsA\** or *ΔzipA ftsA\** cells. Indeed, HK629 Kil’s cell division inhibition activity was suppressed in *ΔzipA ftsA*\* cells (Fig. 5). This strongly suggests that ZipA is also required for the effect of HK629 Kil’ on FtsZ assembly and subsequent bacterial cytokinesis.

**Figure 5.**
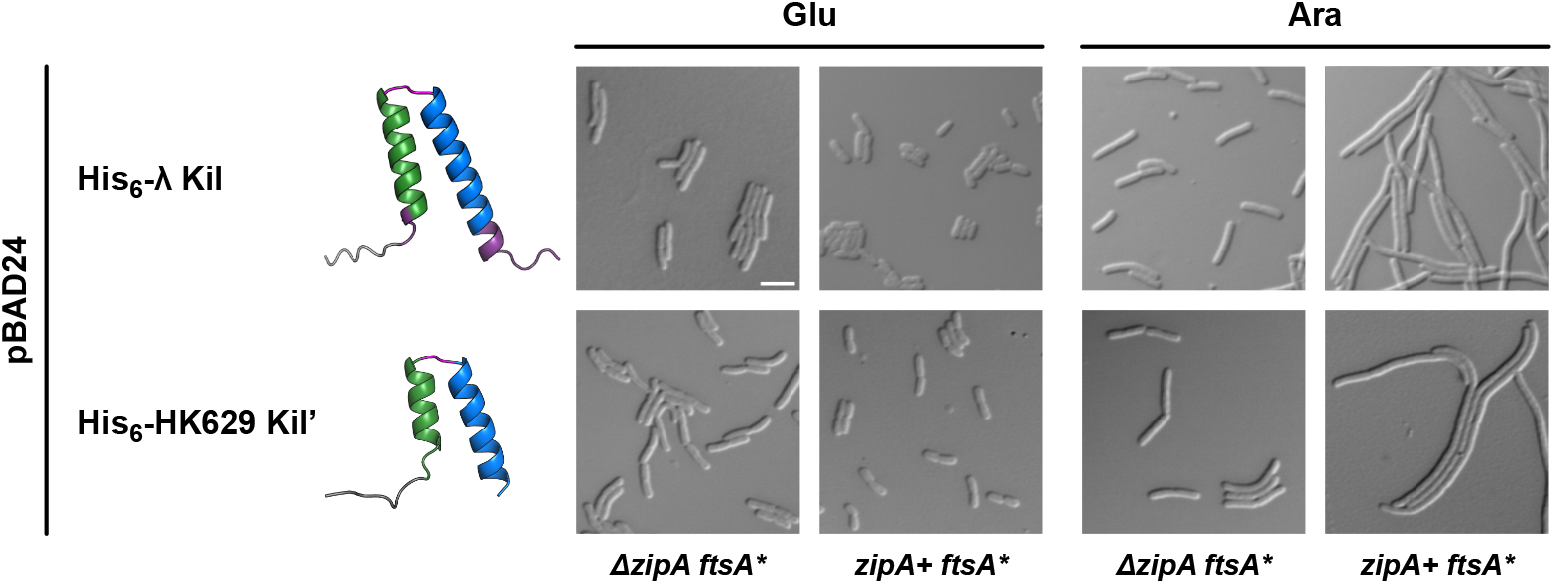
HK629 Kil requires ZipA for its cell division inhibition activity. DIC microscopic images of *E. coli* WM1657 *(ΔzipA ftsA*\*) or WM1659 (*zipA+ ftsA*\*) expressing His-tagged FL λ Kil or HK629 Kil’ from pBAD24 under repressing (0.2% glucose, Glu) or inducing (0.2% arabinose, Ara) conditions. Cells harboring the FtsA^R286W^ variant (FtsA*) that allows bypass of *zipA* (*ΔzipA::aph*) are resistant to the inhibitory effects of λ Kil or HK629 Kil, but are not resistant when containing FtsA^R286W^ in a *zipA*+ background. Ribbon diagrams of both the Kil peptides are shown as modeled by AlphaFold 3. Scale bar, 5 µm.

### 2D. Kil inhibits cytokinesis of UPEC strain

UPEC strains are the primary causative agents of urinary tract infections (UTIs), posing a significant health concern in the United States [32]. The UTI89 strain is widely used to study UPEC physiology and virulence during UTI pathogenesis. To investigate the potential of Kil peptides as an alternative to traditional antibiotics, we introduced pBAD24-His-Kil or -His-HK629 Kil’ into UTI89 and induced expression of the *kil* genes with arabinose. This resulted in the inhibition of cell division and filamentation in bacterial cells expressing His-tagged full-length (FL) Kil or HK629 peptides (Fig. 6). In contrast, *E. coli* cells carrying an empty plasmid showed no change in cell length. These findings suggest that Kil peptides can be potent cell division disruptors in pathogenic strains of *E. coli*.

**Figure 6.**
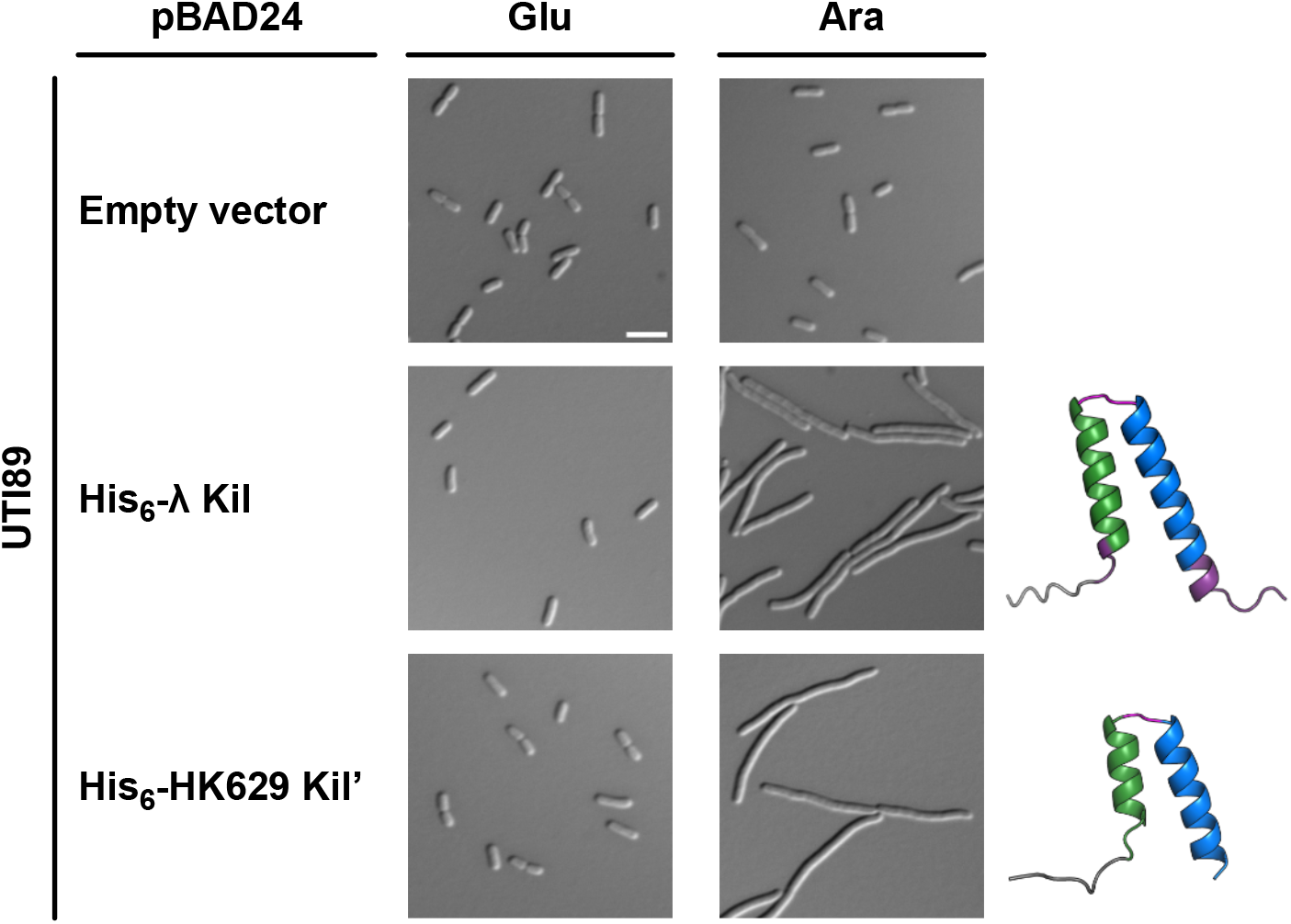
Kil peptides produced from phages HK629 or λ inhibit cell division of Uropathogenic *E. coli* (UPEC) strain UTI89. DIC microscopic images of *E. coli* cells expressing His-tagged FL λ Kil or HK629 Kil’ from pBAD24 under repressing (0.2% glucose, Glu) or inducing (0.2% arabinose, Ara) conditions. Ribbon diagrams of both Kil peptides are shown as modeled by AlphaFold 3. Scale bar, 5 µm.

## 3. Discussion

In this study, we explored the structural and functional features of bacteriophage-derived Kil peptides and assessed their inhibitory effect on cell division of both *E. coli* K-12 and a pathogenic UPEC strain. Structural analysis of the λ Kil peptide revealed a conserved HTH motif essential for its function. Both FL Kil and truncated variants that include this motif were able to disrupt Z-ring assembly, underscoring the critical role of the HTH in retaining Kil’s anti-FtsZ activity. Deletions of the N-terminal five residues or the C-terminal 11 residues at the ends of the two helices did not compromise Kil’s inhibitory function, whereas deleting both simultaneously inactivated Kil’s activity. This suggests that when the 31-residue KilΔN5ΔC11 is combined with either the N-terminal five amino acids or the C-terminal eleven amino acids, it forms the minimal active λ Kil domain which retains the ability of the FL peptide to inhibit *E. coli* cytokinesis. In addition, an intact HTH motif within the minimally active Kil is indispensable. Based on the requirement of ZipA for Kil’s inhibition of FtsZ, we speculate that the two helices in Kil’s HTH may each target ZipA and FtsZ independently. Such a mechanism would result in a two-pronged inhibition of cell division by disrupting both FtsZ polymer dynamics and ZipA’s functions in FtsZ membrane tethering and divisome assembly. A similar two-pronged inhibitory mechanism involves disruption of both FtsZ and MreB function by the *E. coli* CbtA toxin [33]. Understanding how Kil acts on FtsZ and ZipA may shed further light on how FtsZ normally interacts with ZipA and other proteins during bacterial cell division.

The structural and functional conservation of Kil peptides across different bacteriophages further suggests a common mechanism of action. In support of this, the Kil peptide from Enterobacteria phage HK629 exhibited similar cell division inhibition as λ Kil, consistent with its similar HTH fold. HK629 Kil’ is the smallest Kil that still retains function in our experiments. The key difference between HK629 Kil’ and the inactive Kil ΔN5ΔC11 mutant are the charges on the C-terminus of the peptides. Despite its significantly shorter length, HK629 Kil’ retains a C-terminal RSK motif resembling a similar RSSK motif found in the C-terminus of full-length λ Kil. Removal of this motif from λ Kil in combination with the N-terminal truncation in the λ Kil ΔN5ΔC11mutant may be responsible for full inactivation of λ Kil. However, this hypothesis has to be validated in the future, and further experiments will be required to unveil the mechanistic details of the *E. coli* cell division inhibition by HK629 Kil’.

The cross-species activity of Kil highlights the potential of Kil peptides as antibacterial agents. In the context of UPEC, a major causative agent of UTIs, induction of filamentation and inhibition of cell division in UTI89 cells expressing His-tagged FL Kil or HK629 peptides confirmed their efficacy in targeting and disrupting the bacterial division machinery. This finding is particularly pertinent given the rising concern of antimicrobial resistance (AMR) among UPEC strains. The effect of bacteriophage encoded peptides on FtsZ localization during infection by pathogenic *E. coli* was also recently investigated in a neonatal meningitis model system (22).

In summary, our study provides evidence for the efficacy of phage-derived Kil peptides as alternative antimicrobial agents. Their structural conservation, and ability to inhibit cell division in pathogenic bacteria underscore their therapeutic potential. It would be interesting to see if the Kil peptides are active against other bacterial pathogens including *Mycobacterium tuberculosis* where both FtsA and ZipA are absent and SepF is the essential membrane tether for FtsZ (7). Future research should focus on elucidating the precise molecular mechanisms of Kil peptide action and optimizing their stability and delivery.

## Materials and methods

### Bacterial Strains and Growth Conditions

All bacterial strains used in this study are listed in Table 1. Competent cells of XL1-Blue, UTI89, WM4952, XTL937, WM1657, and WM1659 strains were prepared from Luria-Bertani (LB) broths supplemented with 25% (v/v) glycerol stored at -80°C. Recombinant *E. coli* strains ANWM1-ANWM34 were prepared by transforming WM1074, UTI89, and the genetically engineered strains WM4952, XTL937, WM1657, and WM1659 with pBAD24/pET15b-derived plasmids producing the full length (FL) Kil peptide or the deleted variants. Microscopic experiments were performed in triplicate using individual colonies isolated from fresh transformation plates. For cultivation, strains were grown in LB medium with vigorous aeration (200 rpm) at either 30°C or 37°C, as specified. Appropriate antibiotics were added as follows: carbenicillin (100 µg/ml), kanamycin (50 µg/ml), or chloramphenicol (15 µg/ml). Gene expression was regulated using IPTG (isopropyl-β-D-1-thiogalactopyranoside), 0.2% D-(+)-Glucose, and 0.2% L-(+)-arabinose (Sigma-Aldrich).

**Table 1:**
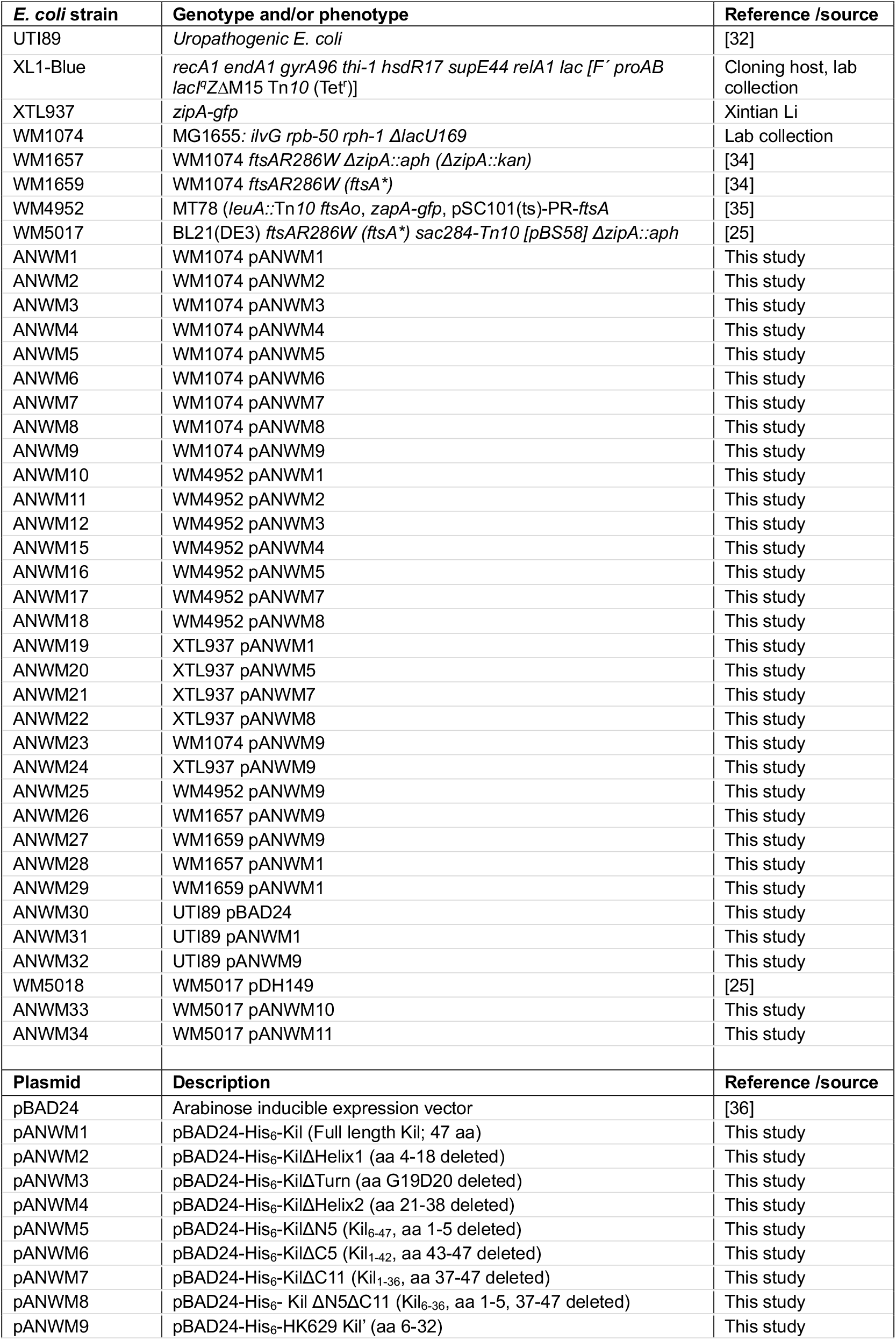
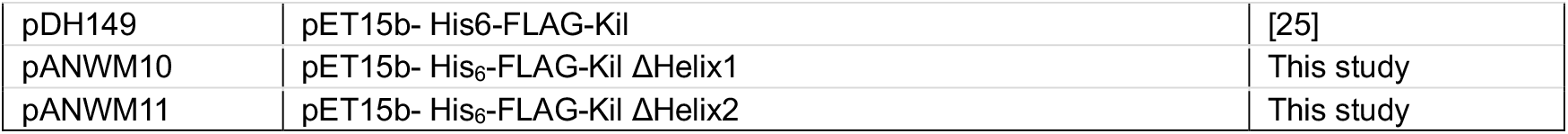
Bacterial strains and plasmids used in this study.

For IPTG inductions of pET15b vector-expressed Kil derivatives, overnight bacterial cultures were diluted 1:100 in fresh LB medium and grown to an optical density at 600 nm (OD600) of 0.7. Protein expression was induced by the addition of 1 mM IPTG, and cultures were incubated for four hours at 30°C. Bacterial cells were harvested by spinning at 7,000 rpm. Supernatants were carefully removed and cell pellets were washed in buffer (50 mM sodium phosphate pH 8.0, 300 mM NaCl), re-centrifuged, and stored at −80°C until used for follow up experiments. For arabinose-inducible protein expression, bacteria were initially grown overnight in LB medium containing 0.2% glucose (Glu, repressing condition). The overnight cultures were then diluted 100 times in 1 ml of pre-warmed LB+Glu. Once the OD600 value reached ∼0.35, the cultures were split into two 500 µl aliquots, pelleted, washed with prewarmed LB, and resuspended in LB with 0.2% arabinose (Ara, inducing condition) and Glu. Following addition of inducers, cultures were further grown for 40 minutes, then observed under light and fluorescence microscopy.

### Plasmid Construction

All plasmids and primers used in this study are described in Table 1 and Table 2, respectively. Recombinant plasmids were constructed using standard molecular cloning techniques. Site-directed mutagenesis (SDM) was employed to generate Kil HTH truncations and shorter deletion mutants on pBAD24 using the templates and primers specified in Table 2. All constructs were verified by DNA sequencing using plasmid specific primers.

**Table 2:**
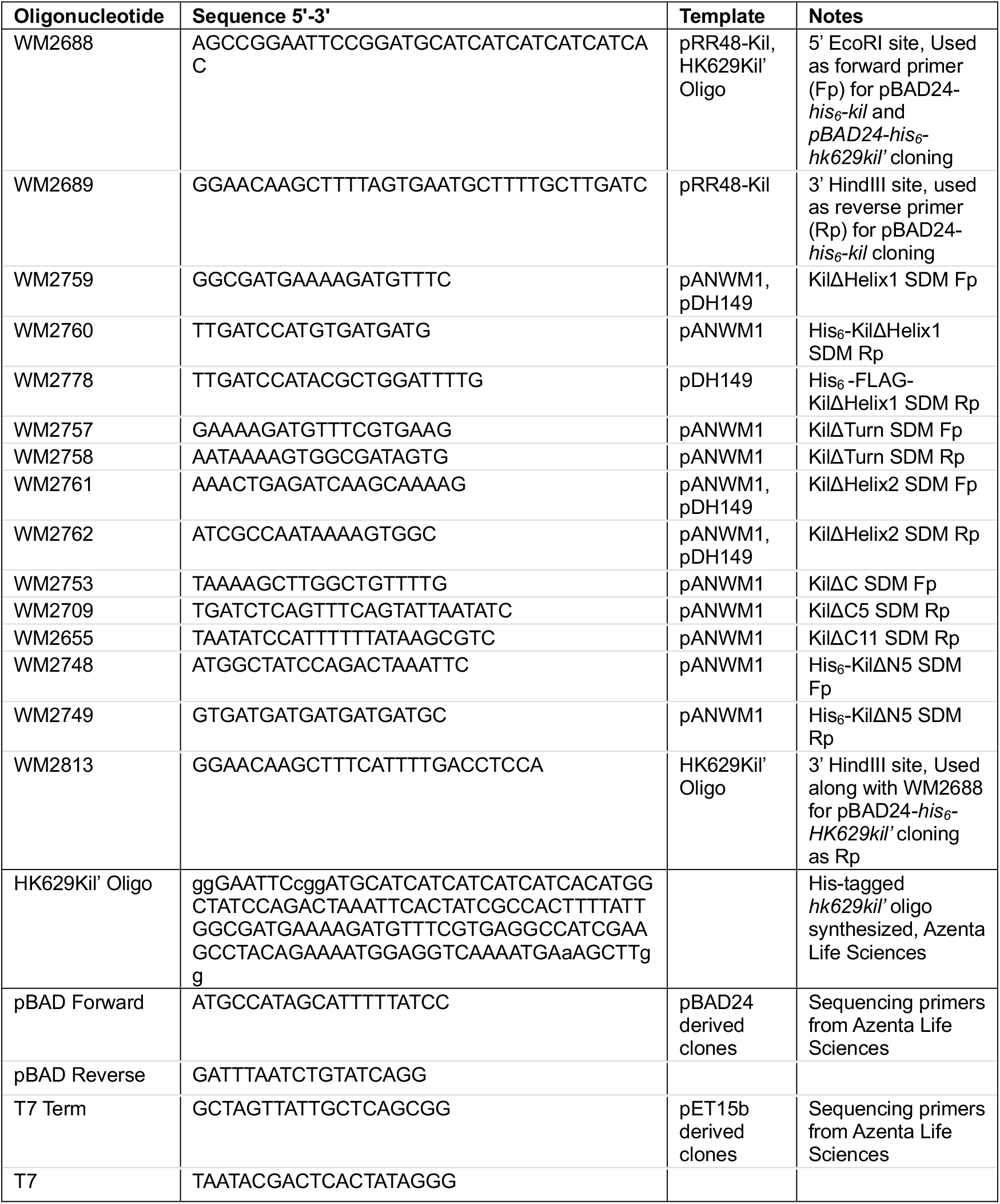
Oligonucleotides used in this study.

### Microscopy

Cells were mounted on agarose pads and imaged using an Olympus BX63 microscope equipped with a Hamamatsu C11440 ORCA-Spark digital complementary metal oxide semiconductor (CMOS) camera and cellSens software (Olympus). Image analysis was performed using Fiji/ImageJ software.

### Immunoblot analysis

To prepare whole cell lysates, bacterial cell pellets were suspended in 1X SDS-PAGE sample buffer. Samples were boiled for 10 min prior to separation by SDS-polyacrylamide gel electrophoresis (PAGE) using 4–20% precast polyacrylamide gels (Bio-Rad, #4561094) with Laemmli running buffer. Spectra Multicolor Low Range Protein Ladder (Thermo Scientific 26628) was used as the molecular weight standard. Samples were transferred to nitrocellulose membranes (Bio-Rad 1620112) using a Trans-Blot SD semi-dry transfer cell (Bio-Rad) at 15 V for 25 minutes min in 1X Tris-Glycine transfer buffer (25 mM Tris, 192 mM glycine, pH 8.3, 20% methanol). Blots were stained with Ponceau stain for assessment of total protein levels, then blocked using 4% bovine serum albumin (BSA) in Tris-buffered saline with Tween 20 (TBST) and incubated with mouse anti-FLAG antibodies (1:5000, Millipore Sigma F1804). After washes, blots were incubated with goat anti-mouse antibody labeled with horseradish peroxidase (1:5,000, Bio-Rad 1706516), developed with Clarity ECL Western Blotting Substrates (Bio-Rad 1705061), and imaged using a ChemiDoc MP imaging system (Bio-Rad).

## ACKNOWLEDGEMENTS

This work was supported by the National Institutes of Health (NIH) Grant AI171856. Part of this work was presented at the annual meeting of the American Society for Microbiology (ASM Microbe 2023), and at the Gulf Coast Consortia AMR conference (GCC-AMR 2024). The authors thank Dr. Jennifer N. Walker for providing the UTI89 strain, Xintian Li for providing the XTL937 strain, and Dr. Xu Wang and Dr. Maria A. Schumacher for useful insights into Kil structures.

## AUTHOR CONTRIBUTIONS

Arindam Naha, Conceptualization, Formal analysis, Investigation, Methodology, Validation, Visualization, Writing – original draft, Writing – review and editing | Todd Cameron, Formal analysis, Methodology, Validation, Writing – review and editing | William Margolin, Conceptualization, Data curation, Formal analysis, Funding acquisition, Investigation, Project administration, Resources, Supervision, Validation, Writing – original draft, Writing – review and editing

