## Supplementary material for "Bacteriophage Kil peptide folds into a predicted helix-turn-helix structure to disrupt *Escherichia coli* cell division": Figure S1

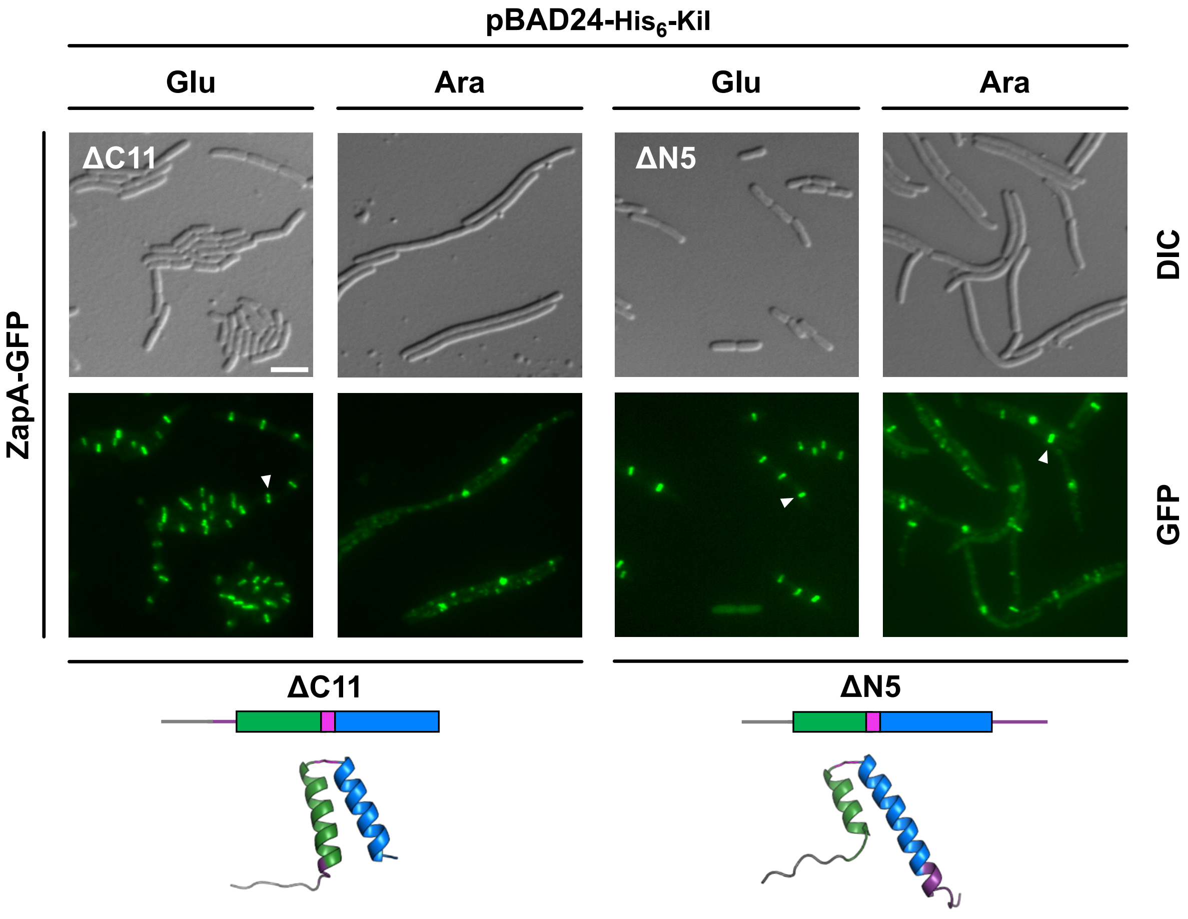


**Figure S1. Shorter λ Kil truncations also inhibit *E. coli* cytokinesis.** (A) Fluorescence microscopic images of WM4952 expressing ZapA-GFP at its native chromosomal locus along with His-tagged KilΔN5 or ΔC11 truncations from pBAD24 under repressing (0.2% glucose, Glu) or inducing (0.2% arabinose, Ara) conditions. Arrowheads indicate intact Z-rings. Schematic diagrams of the Kil derivatives are presented at the bottom of each corresponding micrograph, along with AlphaFold 3-modeled structures. Scale bar, 5 μm.
